# The Evolutionary Genomic Dynamics of Peruvians Before, During, and After the Inca Empire

**DOI:** 10.1101/219808

**Authors:** Daniel N. Harris, Wei Song, Amol C. Shetty, Kelly Lavano, Omar Cáceres, Carlos Padilla, Víctor Borda, David Tarazona, Omar Trujillo, Cesar Sanchez, Michael D. Kessler, Marco Galarza, Silvia Capristano, Harrison Montejo, Pedro O. Flores-Villanueva, Eduardo Tarazona-Santos, Timothy D. O’Connor, Heinner Guio

**Affiliations:** Institute for Genome Sciences, University of Maryland School of Medicine, Baltimore, MD; Department of Medicine, University of Maryland School of Medicine, Baltimore, MD; Program in Personalized and Genomic Medicine, University of Maryland School of Medicine, Baltimore, MD; Laboratorio de Biotecnología y Biología Molecular, Instituto Nacional de Salud, Lima, Perú; Departamento de Biologia Geral, Instituto de Ciências Biológicas, Universidade Federal de Minas Gerais, Belo Horizonte, Brazil.; Centro Nacional de Salud Intercultural, Instituto Nacional de Salud, Lima, Perú

## Abstract

Native Americans from the Amazon, Andes, and coast regions of South America have a rich cultural heritage, but have been genetically understudied leading to gaps in our knowledge of their genomic architecture and demographic history. Here, we sequenced 150 high-coverage and genotyped 130 genomes from Native American and mestizo populations in Peru. A majority of our samples possess greater than 90% Native American ancestry and demographic modeling reveals, consistent with a rapid peopling model of the Americas, that most of Peru was peopled approximately 12,000 years ago. While the Native American populations possessed distinct ancestral divisions, the mestizo groups were admixtures of multiple Native American communities which occurred before and during the Inca Empire. The mestizo communities also show Spanish introgression only after Peruvian Independence. Thus, we present a detailed model of the evolutionary dynamics which impacted the genomes of modern day Peruvians.

---

Native American ancestry is underrepresented by recent whole genome studies^1–4^. As a result, there are numerous questions that remain in both Native American genetic architecture and their early history. One such question regards the peopling of the Americas, which began when the genetically diverse Native American ancestral population^5,6^ diverged from East Asians ∼23,000 years ago (ya)^7^. Following the separation from Asian populations, Native Americans’ entered a period of isolation, potentially in Beringia^8^, for as long as 10,000 years^7,9^. Thereafter, they migrated into the New World, likely through a coastal route^10^, and quickly populated both North and South America. This rapid peopling of the Americas is evidenced by Monte Verde, one of the oldest archeological sites in the Americas, being in Southern Chile and is ∼14,000 years old^11^. It took Native Americans only 1,000 to 2,000 years after the isolation period to populate the majority of the Americas. While it is widely supported that this was a rapid process, questions still remain regarding the divergence between the Amazon, Andes, and coast regions of South America^9^. Peru contains all three regions within its borders and its population has predominantly Native American ancestry^12^, therefore making it an excellent country to study this process.

Peruvian populations have a rich cultural heritage that derives from thousands of years of New World prehistory, that culminated in the last major pre-Columbian civilization, the Inca Empire^13^. The populations of this region have undergone drastic demographic changes that derive from experiences such as forced migration^13–17^ and major population size reduction due to the Spanish conquest, during the 16^th^ century, which introduced mass pandemics to the region^3,4,18–21^. Due to the Spanish conquest, admixture occurred between Native Americans and individuals with European and African ancestry which resulted in modern mestizo populations in addition to Native American ones. These events, along with the original peopling of the region, created a dynamic pattern of evolutionary history and can now be investigated by genomics^1–4^.

To reconstruct the genetic history of Peruvian populations and address the unclear genomic questions about Native Americans, we analyze high coverage whole genome sequence data (N=150) and genotype array data (N=130) to create a geographically diverse dataset representative of Peru (Fig. 1; Supplementary Table 1). Using this data, we show that the Peruvian area was originally peopled ∼11,684-12,915 ya, and that modern mestizo populations originated from multiple Native American sources in addition to their later African and European admixture. We demonstrate that much of this Native American admixture took place before Peruvian independence, and that there has been subsequent admixture between Old and New World populations. These findings establish detailed models of mestizo and Native American evolution in the Andean region, and provide insights into the sociopolitical impacts of genetic variation on the populations of Peru. In providing the largest high coverage genomic dataset of Native American haplotypes to date, this work lays the foundation for understanding the evolutionary history of the Andean region, and the genomic medical needs for this group, and their mestizo cousins^4,22,23^.

**Figure 1.**
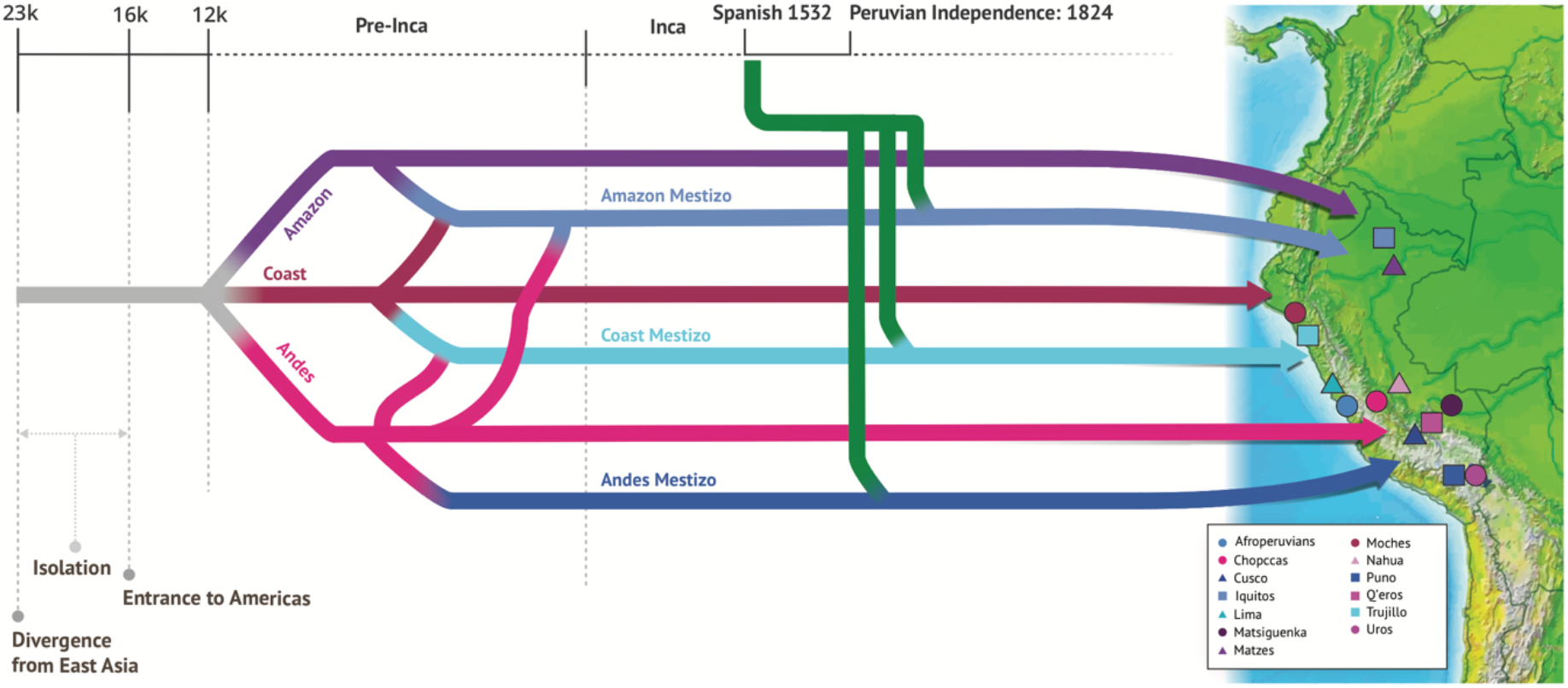
Population history of Peru. No later than 23,000 ya the Native American ancestral population diverged from East Asia and entered a period of isolation^7^. Then ∼16,000 ya the ancestral population began to populate the Americas^7^. We inferred that the studied populations of the three geographic regions in Peru (Amazon, Andes, and Coast) diverged from each other ∼12,000 ya. Prior to the Inca Empire, the Native American populations admixed with each other to form separate populations that would go on to form the mestizo populations. Migration did occur between all regions however we only represent the predominate asymmetrical patterns that shaped modern day populations (Andes to other regions and Coast to Amazon). The Spanish conquerors arrived in Peru in 1532, however the majority of Spanish admixture did not occur until Peruvian Independence in 1824. At this time, the Spanish admixture was only with the prior admixed populations of different Native American ancestries to form the modern mestizo populations while the Native American populations remained essentially isolated. The map on the right of the figure gives the sampling location of these populations and how they correspond to the three major areas indicated in the timeline.

## Results

### Ancestry of Peruvian Populations

We studied13 Peruvian populations that consist of either self-identifying Native American or mestizo (i.e. predominantly admixed ancestry) individuals from the Amazon, Andes, and coast (Fig. 1, Supplementary Table 1). In order to perform a broad characterization of our samples’ ancestry, we combined them with other sources of Native American and global genetic variation^2,24^ (Supplementary Table 2). We find seven ADMXITURE clusters, including three Old World continental sources and four Native American groups that represent Amazonian, Andean, Central American, and coastal ancestries (Fig. 2A and Supplementary Fig. 1). Our samples are enriched for Native American ancestry as all sequenced individuals have Native American mitochondrial haplotypes (Supplementary Table 3), and 103 (59 sequenced) of our samples are estimated to have ≥ 99% Native American ancestry. The high frequency of Native American mitochondrial haplotypes suggests that European males were the major source of European admixture with Native Americans, as others have found^25–27^. The only Peruvian populations that have a proportion of the Central American component are in the Amazon (Fig. 2A). This was observed in other Amazonian populations by Homburger et al.^4^ and could represent ancient shared ancestry or a recent migration between Central America and the Amazon. Our data represent the best reference of Native American haplotypes to date, from high coverage whole-genome sequencing and can serve as a resource for the genetic and genomic analysis of populations worldwide with Native American ancestry, such as Latino populations.

**Figure 2.**
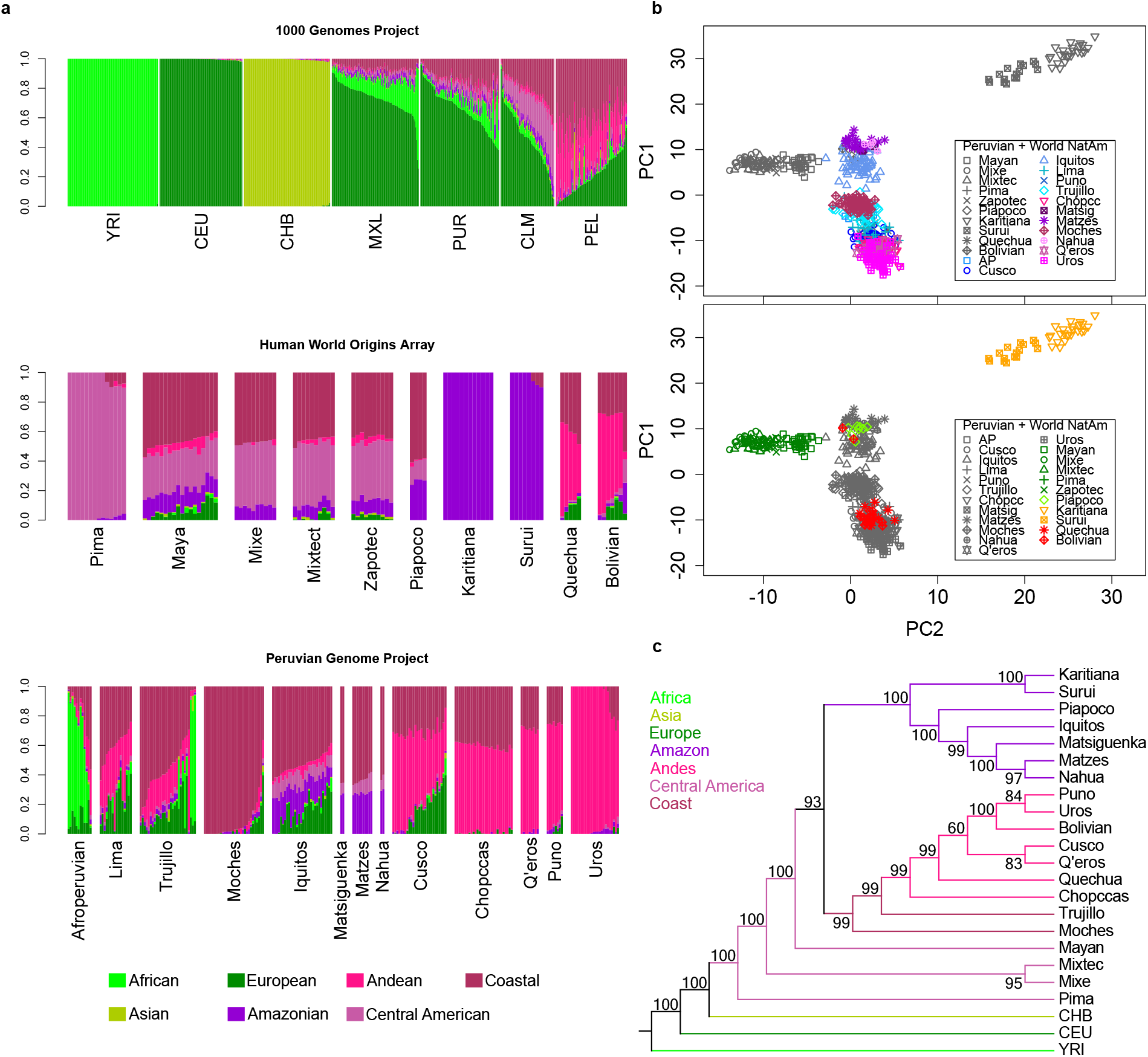
Peruvian population structure. A) Admixture analysis (K=7) of 1000 Genomes Populations, Native American populations from the Human Genome Diversity Panel (HGDP), and all populations from the Peruvian Genome Project. The legend depicts our interpretation of the ancestry represented by each cluster. B) Ancestry specific principal component analysis (ASPCA) of combined Peruvian Genome Project samples with the HGDP. Samples are filtered by their Native American ancestral proportions: ≥ 50%. Each point represents one haplotype. AP and Matsig is short for Afroperuvians and Matsiguenka, respectively. C) A tree made using 1000 Genomes, HGDP, and Peruvian Genome Project samples is shown. Values represent the percent of bootstraps (n=500) in which each node was formed.

The majority of Old World ancestries is European and is mostly seen in mestizo populations. We find little African ancestry in these populations, with the exception of the Afroperuvians and 10 individuals from Trujillo (Fig. 2A). To investigate the Old World sources of European and African ancestry in Peru, we performed Ancestry Specific Principal Component Analysis (ASPCA), which performs a PCA within distinct continental ancestral origins^28,29^ (Supplementary Table 2). As expected for South America based on our understanding of colonial history^4^, the European ancestry component of the Peruvian genomes predominantly comes from Spanish populations (Supplementary Fig. 2), and the African ancestry component predominantly comes from West African populations (Supplementary Fig. 2).

### Biogeography in Peru

To investigate the fine-scale population structure within Peru, evidenced by four Native American ancestry components (Fig. 2A), independent of European and African admixture, we use ASPCA to analyze only the Native American ancestry of individuals’ genomes (Supplementary Table 2). The Native American ASPCA of Native American and mestizo individuals recapitulates the corresponding geographic locations of samples within Peru (Supplementary Fig. 3), as well as the locations of samples within Central and South America (Fig. 2B), which is consistent with previous studies^4,26^. Therefore, there is strong biogeographic signal within the genetic variation of Native American populations as it is possible to determine 6 the approximate geographic location of these populations based on their genetics alone^30,31^. Further, the mestizo populations from the Amazon, Andes, and coast cluster closest to the corresponding Native American populations from their same geographic region, which resembles the same biogeography identified in the Native Americans. (Fig. 2B and Supplementary Fig. 3).

### Founding of Peru

There is a need for better characterization of the peopling of the Peruvian region^9^. As such, we perform a tree-based similarity analysis^32^ of individuals showing ≥ 99% Native American ancestry with other source populations, which reveals clusters within South America that are consistent with our ASPCA and ADMIXTURE results (Fig. 2). We find that the Amazonian cluster is basal to the coastal and Andean populations (Fig. 2C). This suggests an initial peopling of South America by a split migration on both sides of the Andes Mountains (occidental and oriental cordillera). This was likely followed by a subsequent split between coastal and Andean groups. Interestingly, the Moches and Trujillo do not share a common ancestor independent from the Andean populations (Fig. 2C). This is surprising due to the geographic proximity of the Moches and Trujillo (sample sites differ by only 0.01 latitude and 0.002 longitude) (Supplementary Table 1). However, the Moches best represent an un-admixed coastal Native American ancestry, whereas the Trujillo have substantial genetic contributions from both the Andean and coastal Native American populations (Fig. 2A) which likely explains the lack of an independent common ancestor for their entire genome. However we can clearly determine that the separation of coastal and Andean populations from the Amazonian groups occurred first.

To further compare the evolutionary dynamics of Native American populations from these three regions, we construct pairwise population demographic models including effective population sizes and population divergence times^33^ (Supplementary Fig. 4 and Supplementary Fig. 5). We estimate divergence time between regions to be ∼11,684-12,915 ya without gene flow, though we demonstrate through simulation that modeling without gene flow still accurately 7 determines divergence times (Supplementary Fig. 4). The inferred divergence time between the two Andean populations is 10,453 ya, and is statistically distinct from the other population split times (Supplementary Fig. 5). This is also consistent with a model of the rapid peopling of the New World beginning ∼16,000 ya^3,7,34^ because all between region divergence times are not statistically significantly different from each other, when the same Andes population is used in the model (Supplementary Fig. 5).

### Native American Admixture in Mestizo Populations

Following the peopling of the three regions, it appears as the Native American populations remained relatively isolated as the Native American groups tightly cluster within ASPCA and the figure is consistent with geographical location. In contrast, the mestizo populations are found intermediate between the Native American groups, indicative of admixture from multiple Native American groups (Fig. 2B and Supplementary Fig. 3).

To test this more formally, we performed a haplotype-based supervised admixture analysis to further describe this fine-scale structure^35,36^ (Supplementary Fig. 6 and Supplementary Table 4). Results from GLOBETROTTER^35^ illustrate not only the admixture events between New and Old World sources but also the admixture amidst multiple Native American populations (Fig. 3A). Individual mestizo groups show contributions of Native American ancestry from populations in different geographic regions. For example, the Cusco population has greater than 50% of Andean ancestry represented by the Chopccas, Q’eros, and Uros sources along with smaller proportions (<10%) of coastal ancestry from Moches and Amazonian ancestry represented by Nahua, Matsiguenka, and Matzes. Further, the genomic contributions from different Native American sources are correlated to the geographic regions of mestizo genomes. For example, genomic contributions from high altitude populations (Andes) increases gradually from Afroperuvians, Trujillo, Lima and Iquitos in the lowland regions, to Cusco and Puno in the Andes. Similarly, the coastal contribution from Moches decreases from Trujillo to Iquitos, Cusco and Puno, which are far from the coast. Iquitos contains the largest Amazon contribution, from both non-Peruvian and Peruvian Amazon sources with Amazonian contribution lower in coastal and Andean mestizo groups. These results support our hypothesis that mestizo groups have admixture between multiple Native American sources. However, this raises an additional question as to when the admixture between different Native American populations occurred in Peruvian history.

**Figure 3.**
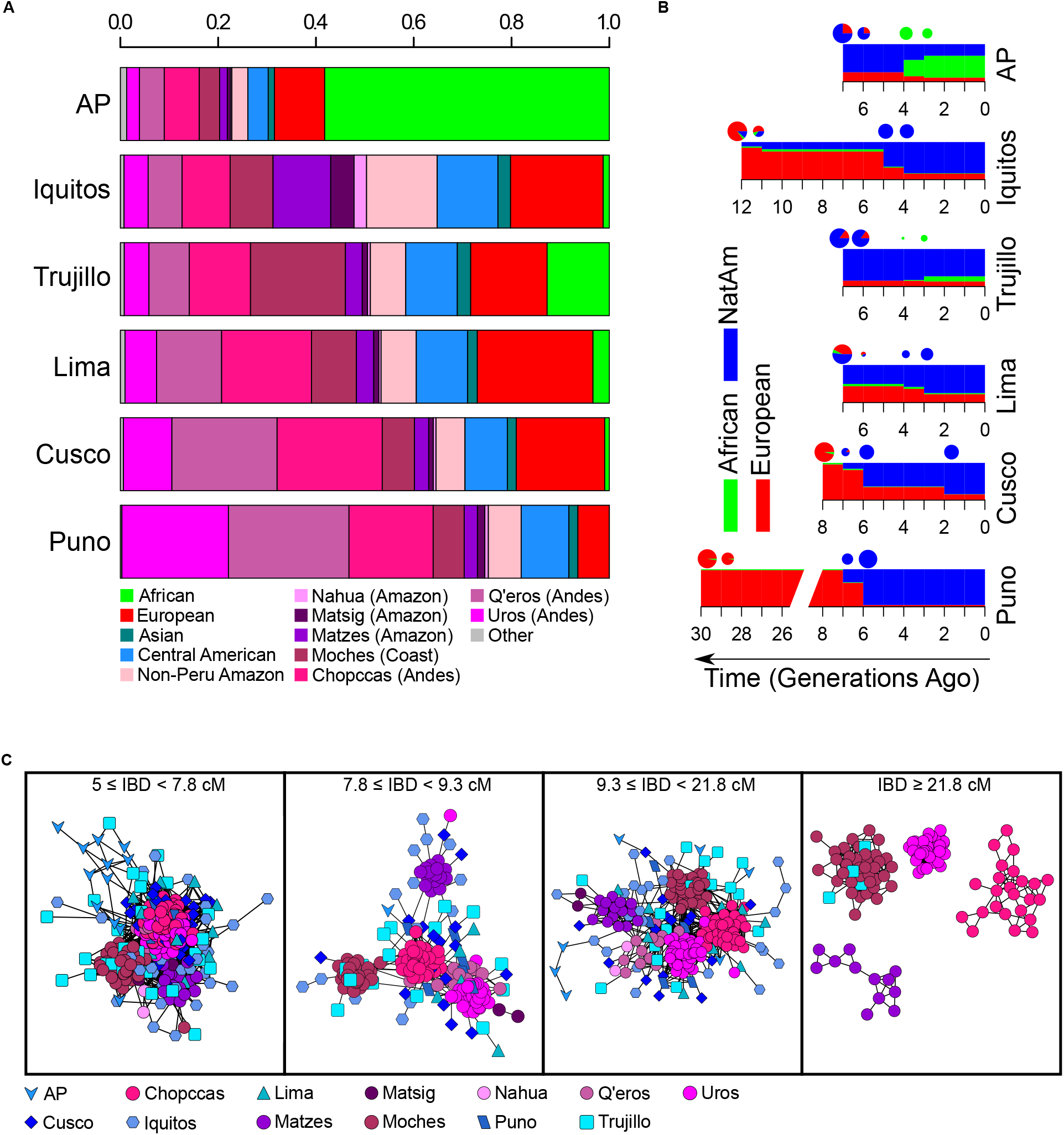
Admixture among Peruvian populations. A) Colors represent contributions from donor populations into the genomes of Peruvian mestizo groups, as estimated by CHROMOPAINTER and GLOBETROTTER. B) Admixture time and proportion for the best fit three-way ancestry TRACT models (European, African and Native American (NatAm)) for six mestizo populations. C) Network of individuals from Peruvian Native American and mestizo groups according to their shared IBD length. Each node is an individual and the length of an edge equals to (1 / total shared IBD). IBD segments with different lengths are summed according to different thresholds representing different times in the past^37^, with 7.8 cM, 9.3 cM, and 21.8 cM roughly representing the start of the Inca Empire, the Spanish conquest and occupation, and Peruvian Independence. IBD networks are generated by Cytoscape^79^. AP and Matsig are short for Afroperuvians and Matsiguenka, respectively.

Therefore, we evaluated the genetic distance between each pair of Peruvian individuals using pairwise Identity-by-Descent (IBD) analysis^37–40^. Since generation time for the most recent common ancestor between two individuals has an inverse relationship to the length of IBD segments shared between their genomes^37,39^, we used the shared IBD segments to infer the individual relatedness at different time periods in Peruvian history^37^. We focus our analysis on the transition between pre-Inca civilizations, the Inca Empire, and Spanish colonial rule based on enrichment of certain lengths of IBD segments, and further combined with their common ancestries across the genome from these time periods (Fig. 3C and Supplementary Fig. 7). Before the Inca Empire (IBD segments of 5-7.8 cM, about 1116-1438 AD), all Andes populations cluster together (Fig. 3C and Supplementary Table 5). During the Inca Empire (IBD segments of 7.8-9.3 cM, about 1438-1532 AD), we identify clear separations between four Native American populations. The Chopccas are at the center of this pattern, and maintain connections directly or through mestizo groups to all other Peruvian populations (Fig. 3C).

During the time period associated with the Viceroyalty of Peru (IBD segments of 9.3-21.8 cM, about 1532-1810 AD), the Native American populations are still tightly connected, but the Chopccas are no longer the major intermediate connector. This is consistent with the historical observation that the location of cultural dominance shifted away from the Andes to the coast and major cities^16^. In addition, mestizo individuals share IBD segments with multiple Native American populations. Therefore, these patterns during and after the time periods of the Inca Empire are 9 consistent with our GLOBETROTTER^35^ results (Fig. 3C and Supplementary Fig. 6), and provide further support that mestizo populations share ancestry from multiple Native American groups. After Peruvian independence (IBD ≥ 21.8 cM, about 1810 AD – present), there were much fewer mestizo individuals sharing IBD segments with Native American groups (Fig. 3C). Therefore, indicating that Native American and mestizo populations became isolated during this time. These IBD results show that mestizo groups’ admixture between Native American populations began before the arrival of the Spanish in 1532.

### Timing European Admixture in Mestizo Populations

In light of these IBD results, we hypothesize that individuals who migrated as a result of dynamics within the Inca Empire and Spanish colonial rule were more likely to later admix with Spanish ancestry individuals. To further investigate this hypothesis we used TRACTS^41^ to estimate the major admixture events between European, African, and Native American ancestries to occur between approximately 1806 and 1866 (Fig. 3B and Supplementary Fig. 8). This suggests that the majority of admixture between the Spanish and Native Americans did not occur until approximately 300 years after Spanish colonization of Peru, which is consistent with what others have found for South America^4^, and may correspond to the sociopolitical shifts of the Peruvian war of independence that occurred between 1810 and 1824. Taken together with our IBD results, this implies that admixture between Native American groups occurred before Peruvian Independence, and that later admixture of these same populations with Europeans led to the modern mestizo populations.

We further tested these observations using Ancestry Specific IBD (ASIBD) segments, which we calculated for all Peruvian samples by intersecting traditional IBD calls with local ancestry inferences (Supplementary Fig. 9). In order to remove the influences from recent shared European and African ancestries, we focused exclusively on Native American genomic components. The ASIBD analysis revealed that mestizo groups are more likely to share IBD segments with multiple Native American populations from different regions across all IBD segment lengths we tested (P-value < 0.001, Supplementary Table 6). We also observe a more recent gene flow into Peruvian mestizo populations from Central America, as these two groups share predominantly large ASIBD segments, which is also consistent with our ADMIXTURE results (Supplementary Fig. 9 and Fig. 2A).

### Genetic Diversity and Clinical Implications for Native American Ancestry Populations

All four Native American populations (Chopccas, Matzes, Moches, and Uros) have small effective population sizes (Supplementary Fig. 5) in comparison to other Native American ancestry populations^3^ and outbred Old World populations^42,43^. This difference could be due to our ability to more robustly estimate demography using only the non-coding regions of the genome obtained by our whole genome sequencing approach. However, these estimates are supported by the observation that Native American populations have a larger proportion of their genome in runs of homozygosity (ROH), as well as decreased heterozygosity counts, compared to mestizo populations (Supplementary Fig. 10 and Supplementary Fig. 11). We also estimated diversity across Peru^44^, and found that Native American populations are in regions of low diversity relative to mestizo populations. Lima (population size = 9,886,647) and Iquitos (population size = 437,376)^45^, two of the largest cities in Peru, have the greatest measures of genetic diversity (Fig. 4A). Further, when Moches are removed from the analysis, the Trujillo join Lima and Iquitos as the most diverse populations in Peru (Supplementary Fig. 12). This signal is robust to the exclusion of known low-diversity populations (i.e. Surui and Karitiana)^46^, to the removal of European admixed individuals, and to single population inclusion/exclusion (Supplementary Fig. 12). These data suggest the coastal and Amazonian cities have greater genetic diversity than the Andean cities.

**Figure 4.**
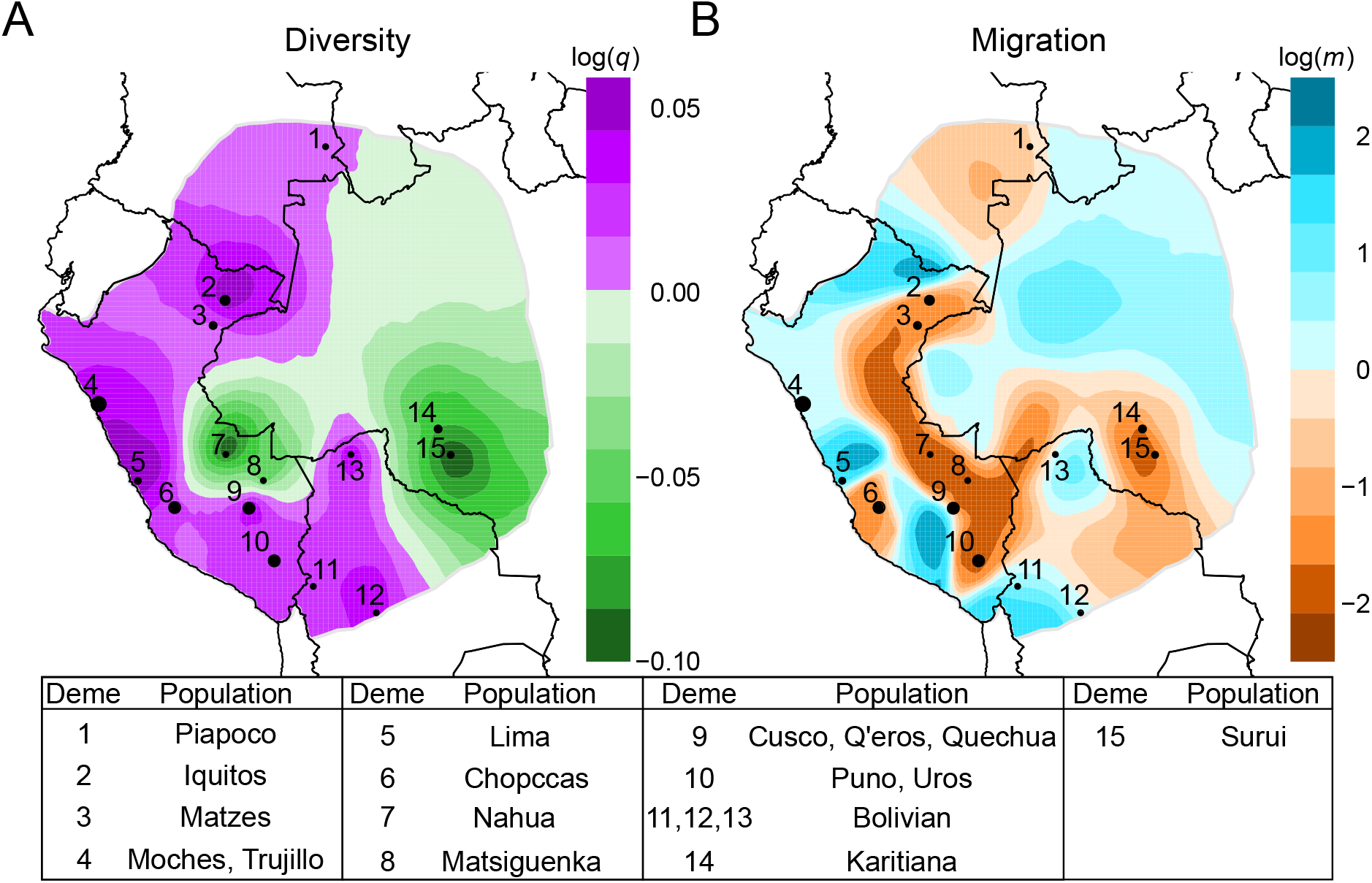
Peruvian Demography. A.) Effective Diversity rates in Peru. Green represents areas of low diversity and purple are high diversity. B.) Contemporary model of effective migration rates in Peru. Brown represents areas of low migration and blue represents areas of high migration. The legend details which populations are grouped into the different numbered demes on the map

The low genetic diversity estimates of Native Americans suggest that there may be an enrichment for rare diseases in Native American ancestry communities^47–49^ living in small populations, as is already observed for isolated populations of different ancestries worldwide^50^. Further, the previously mentioned admixture dynamics in mestizo populations lead us to suspect that they will have greater European specific clinical variation. Due to the underrepresentation of Native American ancestry in genomic databases, we hypothesize that Native American communities may have an increase in recessive disease alleles that are unobserved in current clinical databases^22,51^. In fact, consistent with this hypothesis, we observe fewer ClinVar^52^ variants in our Native American populations than mestizo and Old World populations (Supplementary Fig. 13). Mestizo populations may have inherited risk factors from multiple Native American sources, which further represents the importance of our efforts to better characterize Native Peruvian genomic diversity and disease susceptibility.

### Migration Dynamics in Peru

These low genetic diversity estimates for Native American populations (Supplementary Fig. 10, Supplementary Fig. 11, and Supplementary Fig. 12) and their minimal connectedness in recent IBD networks (Fig. 3C) also suggest that Native American populations are isolated and therefore receive minimal external gene flow. This isolation is further supported by our migration pattern estimates, which show low migration for Native American populations (Fig. 4B), and agrees with prior observations of low genetic diversity (Fig. 4A, Supplementary Fig. 10, and Supplementary Fig. 11). Further, these results are robust to European admixture and individual populations’ exclusions, with two exceptions. First, the Moches exist in an area of high migration, but this changes to an area of low migration when the Trujillo individuals are removed from the model (Supplementary Fig. 14). Therefore, this supports the idea that the Trujillo have received more gene flow from other populations and the Moches have not, which is consistent with our conclusion of low migration and minimal external gene flow for Native American populations (Fig. 2A and Fig. 2C). Second, the removal of Lima does represent a noteworthy complication to this pattern, as a slight migration barrier appears between the Andes and the coast (Supplementary Fig. 14). This suggests that Lima is a crucial intermediate for the migration seen between the coastal regions and the Andes.

As our migration analysis is unable to indicate directionality of migration^44^, we developed an alternative approach based on fine-scale ancestry estimates among the mestizo populations to give the ratio of genomic contributions originating from another geographic region into the local region vs the opposite (Fig. 3A and Supplementary Table 7). We find the indication of asymmetrical migration from the Andes to the coast where the proportion of Andes ancestry which exists in coastal mestizo individuals is 4.6 times more than the coastal ancestry that exists in Andes mestizo individuals (p-value=0.038). Other migrations are from the Andes to the Amazon (ratio=6.3, p-value=0.001), and from the coast to the Amazon (ratio=2.2, p-value=0.002). These results suggest that the majority of migration, at least in terms of mestizo individuals, is in descent of the Andes Mountains toward the cities in the Amazon and the coast.

## Discussion

The Peruvian Genome Project presents high quality WGS and array data with the most extensive sampling of Native American genetic variation to date. As a result, we are able to more accurately model South American demographic history, especially in Peru. We find that the initial peopling of Peru was likely a rapid process which began about 12,000 ya. Following the initial peopling process, there is evidence that mestizo populations received gene flow between multiple Native American sources which likely began before the Spanish arrival and conquest. The majority of this gene flow appears to be in descent of the Andes Mountains which could be attributed to the Inca Empire and/or high altitude selection pressures. The Spanish conquest not only assisted with this admixture between multiple Native American sources, but also resulted in gene flow between the Spanish and these cosmopolitan groups which resulted in modern mestizo populations. Interestingly though, Spanish admixture did not likely occur until Peruvian independence which is ∼300 years after the Spanish first arrived in Peru (Fig. 1).

Since the divergence times between all geographic regions are not statistically different (Supplementary Fig. 5), we cannot confirm the peopling model suggested through tree analysis (Fig. 2C). Therefore, we propose this model as support for one potential scenario of the peopling process where we are confident the first split separated the Amazon populations from the rest of South America. This was then followed by the divergence of the coast and Andes populations. In addition, this implies that the three regions were peopled rapidly as has been suggested for other Native American populations^3,34^. However we find that the separation between these populations is similar to prior estimates of divergence times between Central American and South American populations (12,219 ya), Central American and Caribbean populations (11,727 ya)^3^, and North and South American populations (∼13,000 ya)^7^. Further, this time range also corresponds to early archeological sites in Peru as Monte Alegre from the Amazon, Guitarrero Cave in the Andes, and Amatope on the coast are around as old as 11,000-12,000 years^53^.

These estimates for the peopling process predate the appearance of agriculture in the Amazon by roughly 6,000 years and in the coast by roughly 4,000 years^53^. Furthermore, the divergence between Chopccas and Uros predates the appearance of agriculture in the Andes by ∼400 years^53^. Therefore, it is likely that agriculture was developed after each major geographic region of Peru had been populated for multiple generations. In order to further refine this model, we need denser sampling of these regions and methods that are sensitive enough to distinguish small differences in divergence times.

Following the peopling of Peru, we find a complex history of admixture between Native American populations from multiple geographic regions (Fig. 2B, Fig. 3A, and Fig. 3C). This is likely due to forced migrations by the Inca Empire, known as ‘mitma’ (*mitmay* in Quechua)^13,16,54^, which moved large numbers of individuals in order to incorporate them into the Inca Empire. We can clearly see the influence of the Inca through IBD sharing where the center of dominance in Peru is in the Andes during the Inca Empire (Fig. 3C). A similar policy of large scale consolidation of multiple Native American populations was continued during Spanish rule through their program of “reducciones” or reductions^16,17^. The result of these movements of people created early New World cosmopolitan communities with genetic diversity from the Andes, Amazon, and coast regions as is evidenced by mestizo populations’ ancestry proportions (Fig. 3A). Then, following Peruvian Independence, these cosmopolitan populations were those same ones that predominantly admixed with the Spanish (Fig. 3B). This therefore represents an interesting dynamic where the Inca Empire and Spanish colonial rule created these diverse populations as a result of admixture between multiple Native American ancestries, which would then go on to become the modern mestizo populations by admixing with the Spanish after Peruvian Independence.

These IBD results represent recent migration patterns within Peru centered on the Inca Empire and Spanish Conquerors. However, we also wanted to obtain an overall topography of migration in Peru that is representative of all time ranges since the initial peopling of the region. This migration topography shows that there is a corridor of high migration connecting the city centers in the Amazon, Andes, and coast (Fig. 4B). We also note that there is asymmetrical migration between these regions with the majority of migration being in descent of the Andes towards cities in the Amazon and coast. We therefore hypothesize that this migration pattern may be due to selection pressures for alleles that assist in high altitude adaptation creating a disadvantage to new migrants^55–57^. However, this also aligns with Spanish efforts to assimilate Native American populations^16,17^ and the complex history of Native American admixture in mestizo populations (Fig. 3A and Fig. 3C) which could be a result of Inca Empire policies^13,16,54^. Therefore, future research is required to determine if this migration pattern began during Native American empires or during Spanish rule as compared to being constant since the initial peopling of the region and explore the possibility that this pattern is a result of a combination between the two forces.

These presented evolutionary dynamics of Peruvian people provides insights into the genetic public health of Native American and mestizo populations. Due to the overall low genetic diversity of both Native American and mestizo populations (Supplementary Fig. 10 and Supplementary Fig. 11), it is likely that there is an enrichment of rare genetic diseases unique to these groups. While this is a pattern observed worldwide and has been extensively documented in isolated Old World populations^50^, the diseases that may increase in frequency as the product of these phenomena of isolation are likely to be population specific and should be systematically documented and addressed by the Peruvian health system based on other Latin American experiences^58^ and can add to the growing number of studies finding new insights into biology through studies of isolated populations. Further, the finding that few ClinVar^52^ variants are present in Native American ancestry populations also stresses that these populations are underrepresented in genome studies and require more sequencing efforts. Additional studies will lead to an increase in our understanding of Native American genetic variation and the sequences we present will serve as a reference for future studies. However, we also find strong population structure between the different regions in Peru (Fig. 2B and Fig. 2C), and as a result of the rapid peopling of the New World we hypothesize larger differences in the distribution of pathogenic variants from one group of Native American ancestry to another^3^. This means that future studies must sequence from different regions of the Americas as the populations from other countries in the Americas are expected to be less closely related to these samples. The Peruvian samples we present here likely capture most of the Native American common genetic variation^2^, however large sample sizes in other geographic regions are required to discover rare genetic variation for any given region, which is crucial for understanding genetic causes of diseases, both complex and Mendelian^59,60^.

In conclusion, we present here the largest collection of Native American sequences to date, which makes important strides in addressing the genetic underrepresentation of these populations. In addition, these data help address important questions in Native American evolutionary history as we find that populations from the Andes, Amazon, and coast diverged rapidly ∼12,000 ya. Following the initial peopling of each region, we demonstrate that the majority of the migration in Peru is in descent of the Andes towards the Amazon and Coast. As part of this migration dynamic, we demonstrate that mestizo populations have a complex history of Native American ancestry which occurred prior to their admixture with European ancestry populations following Peruvian independence. The understanding of this complex evolutionary history, in addition to the low genetic diversity of these populations, is crucial for bringing the era of personalized medicine to Native American ancestry populations.

## Online Methods

### Sample Collection

The protocol for this study was approved by The Research and Ethical Committee (OI-003-11 and OI-087-13) of the Instituto Nacional de Salud del Peru. The participants in this study were selected to represent different Peruvian Native American and Mestizo Populations. We obtained informed consent first from the Native American or Mestizo community, and then from each study participant. Native American population cohort participants were recruited from the Matzes, Uros, Afroperuvians, Chopccas, Moches, Q’eros, Nahuas, and Matsiguenka populations. We applied 3 criteria to optimize individuals to best represent the Native American populations: 1) The place of birth of the participant and that of his or her parents and grandparents, 2) their last names (only those corresponding to the region), and 3) Age (eldest to mitigate effects of the last generation). Participants of the mestizo population cohorts were recruited from the cities Iquitos, Puno, Cusco, Trujillo and Lima, and were randomly selected.

### Genomic Data Preparation

150 Native American and mestizo (European and Native American admixed) Peruvian individuals were sequenced to high coverage on the Illumina HiSeq X Ten platform by New York Genome Center (NYGC). Following sequencing, the raw reads were aligned to HG19 and variants were called in each individual’s genome by NYGC. We then used GATK UnifiedGenotyper (GATK), to jointly identify the set of all biallelic single nucleotide polymorphisms (SNPs) that were independently discovered in each individual’s nuclear and mitochondrial genome^61–63^. All SNPs flagged as LowQual or a quality score < 20 were removed^64,65^. We used the webserver of Haplogrep^66–68^ to determine the Mitochondrial haplogroup of all 150 Peruvian WGS samples based on the unified mitochondrial variant database. KING^69^ was used to identify related individuals in our WGS data and we removed the smallest set of individuals to create a final WGS dataset with no pairs having a Kinship coefficient ≥ 0.044.

An additional 130 Native American and mestizo Peruvian individuals were genotyped on a 2.5M Illumina chip. To create a combined Peruvian WGS and array dataset with 280 Native American and mestizo Peruvians, the intersect of the two datasets was filtered to remove all A-T and G-C sites, and any tri-allelic sites created by combining the two datasets that were not due to a strand flip in either dataset. The array-WGS dataset was filtered to remove sites with ≥ 10% missingness or ≤ 2 minor allele count^64,65^. We then ran KING^69^, with default settings, to determine additional relatedness in the merged dataset and to determine the optimal number of individuals with no pair of ≥ 0.044 Kinship coefficient, which consisted of 227 individuals. WGS samples were preferentially chosen in pairs of related individuals between a WGS and array sample. In addition, the WGS individuals were selected from the duplicate samples (ie both array and WGS data for the same sample). A final combined dataset of 4694 samples was created, which contained 227 unrelated samples from Peru, 2504 samples from 1000 Genomes Project^2^ and 1963 Human Genome Diversity Panel (HGDP) samples genotyped on the Human Origin array^24^ by taking the intersection of all three datasets. There were 183,579 markers remaining after removing all A-T, G-C and triallelic sites from the combined dataset^64,65^.

### Admixture Analysis

Using the final combined dataset, we extracted all HGDP Native American individuals genotyped on the Human Origins Array^24^, our samples, and the YRI, CEU, CHB, CLM, MXL, PEL, and PUR 1000 Genomes Project samples^2^. We then applied the same data filters as in the Kinship filtering methods and utilized Admixture v1.23^70^ on K 1-12, using the random seed option, 20 times for each K. We then selected the best log likelihood run for each K and compared Cross Validation (CV) values to determine the K that best fit the data, which in our case was K=7 (Supplementary Fig. 1).

### Phasing and local ancestry inference

The combined dataset was phased using Beagle 4.0^40^ and HapMap Human genome build 37 recombination map^71^. Local ancestry inference was performed using RFmix^72^ to categorize all the markers in the admixed individual genomes according to their similarity to African, European, or Native American ancestry. The European and African ancestries consisted of populations selected from the HGDP genotyped on the Human Origins Array and the Native American ancestry consisted of two parts: 1) populations from the HGDP genotyped on the Human Origins Array (NatAm1) and 2) the Peruvian populations (NatAm2) (Supplementary Table 2). According to the results from ADMIXTURE (Fig. 2A), only samples with ≥ 99% of Native American component were selected as Native American reference haplotypes. As a result, there were 331, 183 and 158 samples used as European, African and Native American references, respectively.

### Ancestry Specific Principal Component Analysis (ASPCA)

We used the approach developed by Browning et. al.^28^ to perform the Ancestry Specific PCA (ASPCA). PCAdmix^73^ was used to prune the markers according to their allele frequency and LD with default thresholds and ASPCA was then performed based on the remaining 100,202 markers. To reduce the noise for ASPCA associated with different ancestries, we only included individuals with ≥ 30% European ancestry, ≥ 10% African ancestry, and ≥ 50% Native American ancestry for the European, African, and Native American ASPCA respectively.

### CHROMOPAINTER and GLOBETROTTER

CHROMOPAINTER^36^ explores ancestry using SNP data of haplotypes sampled from multiple populations and reconstructs recipient individual’s genome as serials of genetic fragments from potential donor individuals. The donor group contained 47 populations, from five ancestries: 1) European, 2) African, 3) Native American (NatAm3), 4) Asian and 5) Oceanic, as listed in Supplementary Table 2. Each selected donor population had > 2 samples to avoid spurious estimation of ancestry (except for Peruvian Matsiguenka and Nahua populations because these populations only have 2 samples and represent variation we are keenly interested in). All six Peruvian mestizo populations are treated as recipient populations. Due to the computational complexity, we estimated the parameters “recombination scaling constant” and “per site mutation rate” using five randomly selected chromosomes (1, 2, 6, 10 and 16) with 10 iterations of expectation-maximization algorithm, as suggested by Montinaro et. al^74^. The two estimated parameters were 225.08 and 0.00073, respectively. We “painted” both the donor and recipient individuals’ genomes” using the combination of fragments from all donor chromosomes. The companion program GLOBETROTTER^35^ was then used to estimate the ancestral contributions from different donor populations into the recipient populations, with DNA information on multiple sampled groups (as summarized by CHROMOPAINTER). In this way, we could achieve finer description of population structure.

### Directional migration for mestizo populations

In order to infer the direction of migration inside Peru, we defined a parameter, *ratio* = 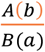, where *A* and *B* are mestizo populations from two different geographic regions, while *a* and *b* correspond to the Native American ancestry sources from these two regions. The three geographic regions are Amazon (represented by Iquitos), Coast (represented by Trujillo and Lima) and Andes (represented by Cusco and Puno), which contains all the mestizo samples in that particular region. *A(b)* means the average proportion of individual’s’ genome originated from region *B* which then existed in individuals living in region *A*. Using only the mestizo populations, we first calculated the original ratio for each combination of regions *A* and *B*. Then we randomly permuted the regional labels for individuals from these two regions and each time a shuffled ratio was calculated by recomputing the population admixture proportion with the new labels. Empirical p-values were then calculated based on comparing the original with the 1000 permutations.

### Normal and Ancestry Specific IBD (ASIBD)

For the phased dataset of Peru with HGDP genotyped on the Human Origins Array, genomic segments that are identical-by-descent (IBD) between individuals were estimated between all pairs of samples on haplotypes with Beagle 4.0^40^. Based on the length of normal IBD segments, an approximation can be used to infer the past generation time^37^: generation ago 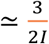, where / is the length of IBD segment in Morgans. Our IBD network analysis are based on all IBD segments that are longer than 5 cM to avoid possible background noise.

For all time frames we had sufficient signal to identify patterns. Supplementary Fig. 7 gives the distribution of the number of segments, with larger segments being fewer, but clearer in terms of the signals they represent, ie. if you have a large segment it is less likely to be a false positive and represent its signal well. For example, during the Inca Empire we use segments of length 7.8 to 9.3 cM which constitute 13% of all segments identified greater than 5 cM.

In addition to normal IBD, we also defined the Ancestry Specific IBD (ASIBD) using the local ancestry segments information from RFMix. If two haplotypes from different individuals share IBD segments and part of the shared segments come from the same ancestry, then the overlapping part between shared IBD and common ancestral segments is defined as Ancestry Specific IBD. In our IBD analysis, to preserve the diversity of Peruvian samples, we kept 259 individuals from Peru with ≥ 50% of Native American component in their genomes, including those related ones. Because the mestizo samples had shorter Native American component in their genomes than the Native American samples, the significance of their Native American IBD might be underestimated. To avoid this, we calculated the total lengths of common Native American segments among all pairs of haplotypes (between different individuals) and used the shortest length as a cap threshold for all other pairs, which was about 700.4 cM between the haplotypes from one Afroperuvian sample and one Cusco sample. For each pair of haplotypes, their common Native American segments were randomly picked without replacement until the accumulated length reached this threshold value. These selected Native American segments were then combined with their shared IBD segments to estimate the Native American specific IBD between the two haplotypes. We could then have equal comparison for all individual pairs with different Native American proportions in their genomes, by using the value of Native American segment length per shared IBD length (NatAm / IBD).

We statistically evaluated the hypothesis that mestizo individuals were more likely to be admixed from multiple Native American populations. We accomplished this by defining four possible population categories, including three Native American based on their geographic origin: Amazon (represented by Matzes, Matsiguenka and Nahua), coast (represented by Moches) and Andes (represented by Chopccas, Q’eros and Uros), along with one mestizo group (represented by Afroperuvians (AP), Iquitos, Cusco, Puno, Lima and Trujillo). Based on the Native American specific IBD network, we defined the connection pattern *N*_1_(*i*) – *M*(*l*) – *N*_2_(*k*) as two distinct individuals, *i* and *k*, who come from two different Native American geographic regions, *N*_1_ and *N*_2_ where *N*_1_≠*N*_2_, and connect to each other through a mestizo individual, *I*. For example, Coast(i)-mestizo(Z)-Andes(/c), *i, I, k* are the sample IDs. Similarly, *N*_1_ *i* – *N*_2_ *j* – *N_3_(k)* represents the pattern of two distinct individuals, *i* and *k*, who come from two different Native American geographic regions, *N_1_* and *N_2_*, connect to each other through an individual *j*, who comes from either the same (if *N*_1_ = *N*_2_ ≠ *N*_3_, eg, Uros(i)-Chopccas(/)-Moches(/c), as both the Uros and Chopccas come from the Andes) or a different Native American label (if *N*_1_ ≠ *N*_2_ ≠ *N*_3_, eg, Moches(*i*)-Chopccas(*j*)-Matzes(*k*), where here *N_1_* is the Coast region, *N_2_* is the Andes region and *N_3_* is the Amazon region). The ratio of number of *NMN* connections to number of *NNN* connections represents the frequency of the mestizo individuals being mixed from different Native American groups. To test for significance, we then randomly permuted the labels for all individuals in an IBD network and each time a shuffled ratio was calculated by re-computing the numbers of two patterns with the new labels. Empirical p-values are based on 1000 permutations.

### TRACTS

TRACTS^41^ was used to estimate the admixture time and proportion in six mestizo populations from Peru admixed by European, African, and Native American ancestries, according to the local ancestry inference by RFmix. Ancestral segments shorter than 11.7 cM were not used for model optimization because their numbers might not be accurately estimated, as suggested by Baharian et. al^37^. For each mestizo population, five models were optimized, including two 2-ancestry and three 3-ancestry models. Then the best-fit models were chosen according to their Bayesian information Criterion (BIC) scores.

For two-ancestry admixture, each model assumes Native American (NatAm) and non-Native American (Non-NatAm, which combines European and African) components. The first model, p_1_p_2_, represents the admixture between different ancestries p_1_ and p_2_ in a single migration event. The second model, p_1_p_2__p_1_x, represents the second incoming migration event from ancestry p_1_ in addition to the initial admixture between p_1_ and p_2_, where x indicates no migration from the other ancestry p_2_. p_1_ and p_2_ could be either one of NatAm and Non-NatAm ancestries.

For three-ancestry admixture models we assumed European, African, and NatAm components (Fig. 3B). The first model, p_1_p_2_x_xxp_3_, represents the initial admixture event between ancestries p_1_ and p_2_, then followed by the second incoming migration event from ancestry p_3_. The second model, p_1_p_2_x_xxp_3__xxp_3_, represents a third migration event from ancestry p_3_ again into the admixed population, in addition to the first p_1_p_2_ and the second p_3_ admixture events. Similarly, the third model, p_1_p_2_x_xxp_3__p_1_xx, represents the third migration event from ancestry p_1_ again in addition to the first and second admixture events. p_1_, p_2_ and p_3_ could be one of European, African, and NatAm ancestries and are different from each other. Again, we used Bayesian information Criterion (BIC) to select the best model, while penalizing for over parametrization.

### Diversity Calculations

We calculated the number of ClinVar variants present in Peruvian and the YRI, CEU, CHB, and PEL from the 1000 Genomes Project using CLNSIG equals to 5 from ClinVar^52^ and we also required that the minor allele frequency was <5%^75^. The median number of variants is ≤ 2 for all Peruvian populations, including PEL from 1000 genomes^2^, as well as the Asian population CHB. The Native American populations have a median value of 1. This is in comparison to the greater numbers of African (YRI) and especially European (CEU) ancestry individuals.

As a measure of inbreeding and diversity, we calculated runs of homozygosity (ROH) in all non-related individuals’ autosomes from the WGS dataset and removed sites with ≤ 2 minor allele count (MAC) and ≥ 10% missing genotypes. We used the plink command -- homozyg to calculate: 1) number of ROH segments, 2) average Size of ROH segments, and 3) total genome content in ROH per individual^64,65^. As an additional metric of genetic diversity, we calculated the number of heterozygous sites per individual from the WGS dataset. The same genotype quality and kinship filters were applied as previously stated. For each individual, a site was considered heterozygous if the most parsimonious genotype call was that of “0/1”, i.e. a call representing a genotype containing one reference allele and one alternate allele.

### Tree Analysis

All individuals identified as being ≥ 99% Native American ancestry, through Admixture analysis (Fig. 2A), and the YRI, CEU, and CHB from the 1000 Genomes Project were extracted from the final combined dataset and applied the same filters as in the kinship analysis. We then constructed a tree using treemix v1 .12^32^ over 500 bootstraps, with the YRI set as the root of the tree and the cluster set to 1 SNP. We used PHYLIP consense tree v3.68^76^ to calculate the consensus of the 500 bootstraps and then used MEGA v7.0.14^77^ to plot the consensus tree.

### δaδi modeling

We calculated the site frequency spectrum, using δaδi^33^, to estimate a simple demographic history of our Native American populations similar to the model implemented by Gravel et al ^3^. We removed the influence of European admixture in our model by selecting individuals from the Chopccas, Matzes, Moches, and Uros populations with ≥ 99% Native American ancestry as calculated by ADMIXTURE analysis (Fig. 2A). Using WGS data only, we removed gene regions from the autosomes of these individuals by excluding all positions within +/- 10,000 bp from all genes in RefSeq downloaded from the UCSC Genome Browser on February 16, 2016^78^. We further removed sites with ≥ 10% missing genotype values, and lacked a high quality human ancestor^2^ trinucleotide (including the variant and +/- 1 bp) which resulted in a total of 2,891,734 SNVs. We then jointly called all positions in the autosome, excluding prior identified variant positions, in all individuals used in δaδi analysis with GATK joint genotyper^61–63^. The same WGS and gene region filters as before were applied and we then added this discovered high quality invariant sites to the total variant sites to yield the entire sequenceable size of our dataset to be 1,025,346,588 base pairs. We then performed all pairwise two-population models among the Chopccas, Matzes, Moches, and Uros calculating the following parameters: 1) N_A_ = ancestral effective population size change, 2,3) N_1_, N_2_ = effective population size of population 1 and 2, 4) T_A_ = time of ancestral effective population size change, and 5) T_S_ = time of divergence between population 1 and population 2 (Supplementary Fig. 5). We down-sampled the Matzes by 2 haplotypes and the other three populations by 4 haplotypes. We utilized the ll_multinom optimizer and corrected the SFS with the following files provided in the δaδi installation^33^: “tri_freq.dat”, “Q.HwangGreen.human.dat”, and “fux_table.dat”. Each model was run 1000 times with the maximum iterations set to 1000000. The run with the maximum log likelihood was selected to best represent the model.

We calculated 95% confidence intervals (CI) by performing 500 bootstraps and removing the upper and lower 2.5% of the inferred values. We first formed 1 MB segments of sequenceable bases across each chromosome (final segment is often truncated as it was not 1MB long). We then formed each bootstrap by randomly selecting, with replacement, these 1MB genomic segments to equal the total sequenceable size of our dataset. Then each δaδi model was run on the bootstraps with the same method as the initial inferred value analysis. We used a human generation time of 30 years and a mutation rate of 1.44 x10-^8^, as calculated for Native American ancestry by Gravel et al.^3^ to convert estimated values into chronological time and effective population sizes.

To assess the accuracy of our demographic model, we simulated 50 replicates of three populations with samples sizes of 7 (population 1), 20 (population 2), and 40 (population 3) (consistent with our own population numbers) with a divergence time of 12,000 years ago (ya). We performed three comparisons with these populations: 1) comparison A is between population 1 and population 2, 2) comparison B is between population 1 and population 3, and 3) comparison C is between population 2 and population 3.

### EEMS migration modeling

To create a contemporary model of migration and diversity in Peru, we used estimated effective migration surfaces (EEMS) to model the effective migration and diversity rates^44^ of all non-related Peruvian Genome Project samples and HGDP samples genotyped on the Human Origins Array^24^ from Peru, Bolivia, Colombia, and Brazil with < 1% African ancestry. We also ran EEMS on individuals with ≥ 99% Native American ancestry and removed the Lima from the original EEMS analysis to reveal any effects European admixture had on our model. We attempted to run EEMS modeling on all individuals with < 99% Native American ancestry, however the resulting migration topographies in each model did not converge even though the MCMC chain did. This is likely due to the filter leaving a small number of individuals and demes which created too many local maxima within the data so it is not possible to discern the optimal < 99% Native American ancestry model. In addition, to further test the robustness of our model we removed the following subsets of populations from the African ancestry filtered dataset: 1) Trujillo, 2) Moches, 3) Puno, 4) Uros, 5) Cusco, 6) Quechua, 7) Cusco and Quechua, 8) Q’eros, 9) Karitiana, 10) Surui, and 11) Karitiana and Surui. In all EEMS runs we removed SNPs with ≥ 10% missingness^64,65^. We optimized EEMS parameters by adjusting the qEffctProposalS2, qSeedsProposalS2, mEffctProposalS2, mSeedsProposalS2, mrateMuProposalS2, and negBiProb such that the acceptance proportions for all parameters, except degrees of freedom, were within 10-40%, as suggested by Petkova et al.^44^ All models were tested with 300 demes and the MCMC chain was run for 15 million iterations with a 14 million iteration burn-in for nine independent runs to insure the MCMC chain converged to the optimal Log posterior. We then plotted the migration rates, diversity rates, MCMC chain log posterior, dissimilarities within sampled demes, dissimilarities between pairs of sampled demes in relation to fitted dissimilarities and geographic distance between demes with the EEMS distributed R scripts^44^. We ensured that at least 3 of the 9 MCMC chains converged and that the dissimilarities within and between demes met the accepted distributions as specified by Petkova et al.^44^ We then replotted all migration and diversity plots for each model with only the MCMC chains that converged to the maximum log posterior probability.

## Acknowledgments

We would like to thank Susan O’Connor, Claire Fraser, Lance Nickel, and Ruth Shady Solis for their constructive comments and perspectives. This work was funded under the Center for Health Related Informatics and Bioimaging at the University of Maryland School of Medicine (D.N.H., M.D.K., A.C.S., and T.D.O.), institutional support for the Institute for Genome Sciences and Program in Personalized Genomic Medicine at the University of Maryland School of Medicine (T.D.O.) and institutional support from the Instituto Nacional de Salud, Lima, Perú (K.L., O.C., C.P., D.T., V.B, O.T., C.S., M.G., H.M., P.F.-V., and H.G.). We would also like to thank all the people that facilitated the recruitment of participants including the following the Direcciones Regionales de Salud from Loreto, Puno, Cusco, La Libertad, Huancavelica, Ica, Piura, Ancash, Arequipa, Ayacucho, Tacna, Ucayali, San Martin, Amazonas, the Universities (Universidad Andina Nestor Caceres Velasquez-Facultad de Ciencias de la Salud, Universidad Nacional Jorge Basadre Grohmann, Universidad Nacional Mayor de San Marcos, Universidad Nacional San Agustín, Universidad Nacional de San Cristóbal de Huamanga, Universidad Nacional Santiago Antúnez de Mayolo and Universidad Nacional de Trujillo, and all the participants in this study.

## Author contributions

T.D.O., H.G., O.C, C.P., conceived of the project. D.N.H., W.S., A.C.S., M.D.K., and T.D.O. performed all population genetics analyses. O.C, C.P and K.L generated the genotype data and V.B. and E.T.-S. performed quality control analyses (QC) on these samples. K.L., O.C., C.P., D.T., O.T., C.S., M.G., S.C., H.M., P.O.F.-V., and H.G. collected and quality controlled the DNA samples. D.N.H, W.S., A.C.S, and T.D.O. sequenced and QC the WGS data from New York Genome Center. All authors contributed to the writing of the paper.

## Competing financial interests

We declare no competing financial interests. 29

